# GEM-GPT Enables Personalized Cell Type-Resolved Therapeutic Design for Systems Pharmacology

**DOI:** 10.64898/2026.07.17.739269

**Authors:** Shuo Zhang, Rahul Ohlan, Mohammadsadeq Mottaqi, Lei Xie

## Abstract

Generative AI is transforming drug discovery, yet most approaches follow one-drug-one-target paradigms ill-suited to the heterogeneity of chronic, systemic diseases. Systems pharmacology offers an alternative, but generative tools designed for it remain scarce. We introduce GEM-GPT, a transcriptomics-guided framework that generates personalized therapeutic candidate molecules intended to shift cell type-specific disease states toward healthy phenotypes. GEM-GPT uses a biology-inspired deep fusion architecture that couples a single-cell RNA-sequencing foundation model with a molecular GPT, modeling cell type-specific chemical-gene interactions throughout molecule generation rather than through fixed conditioning. Across bulk and single-cell chemical perturbations and CRISPR knock-out signatures, GEM-GPT outperforms state-of-the-art baselines, produces cell type-resolved molecules, and generalizes to unseen cellular contexts. In a case study on opioid use disorder (OUD), it generates novel candidates, recovers FDA-approved OUD-related drugs absent from training, and yields predicted binders to OUD-related targets. GEM-GPT bridges single-cell omics and molecular generation for personalized, cell-type-resolved, systems-aware therapeutic design.

## 1 Introduction

Drug discovery and development are among the most resource-intensive and high-risk endeavors in biomedical science. A new drug requires on average 10–15 years and over $1–2 billion to progress from initial concept to regulatory approval [1]. Despite substantial investment in target identification, lead discovery and optimization, and preclinical validation, approximately 90% of drug candidates that enter clinical trials ultimately fail [1, 2]. A major contributor is the continued reliance on the *one-drug-one-target* paradigm [3, 4]. Although this strategy has produced important successes, particularly when disease biology is dominated by an actionable molecular lesion [5], it can oversimplify chronic and systemic disorders that are polygenic, multifactorial, and dependent on cellular and physiological context [3, 4, 6]. In such diseases, modulation of a single gene may be insufficient to restore homeostasis across the broader biological network [7, 8]. Systems pharmacology and network-based drug discovery therefore emphasize coordinated modulation of disease-relevant networks rather than isolated targets [3, 6, 9], yet AI-driven tools that implement these principles for *de novo* therapeutic design remain limited.

An increasingly important goal in drug discovery is the design of therapies tailored to the individual patient. Inter-individual variability in genetic background, epigenetic state, environmental exposure, and tissue-specific gene expression can produce substantial differences in treatment response [10–12]. The past decade has witnessed the rise of precision medicine, wherein patient-specific molecular profiles guide therapeutic decision-making [11, 12]. However, precision drug design, defined here as *de novo* generation of molecules tailored to an individual’s molecular disease state, remains challenging because conventional target-based workflows generally do not encode the cellular context that distinguishes patients.

Single-cell RNA sequencing (scRNA-seq) provides a route to incorporate this context by resolving disease-associated transcriptional programs at cellular resolution [13, 14]. Therapeutic molecules could therefore be designed to shift patient- and cell type-specific disease states toward healthy states without requiring a predefined target, an approach particularly relevant to complex diseases with incompletely understood mechanisms [15, 16]. Nevertheless, compared with target-based drug discovery, relatively few computational methods have been developed for systems pharmacology approaches centered on the modulation of cellular states [17–20], and the integration of single-cell transcriptomics with *de novo* molecular generation remains largely open [20].

The emergence of foundation models pretrained on large-scale biological data has provided new computational tools for addressing this challenge. scRNA-seq foundation models [21–24] have demonstrated the capacity to learn generalizable representations of cellular states, while generative models for molecular design have advanced rapidly [25–29]. Several recent works have begun to bridge transcriptomics and molecular generation. MolGAN [17] conditions a generative adversarial network on bulk transcriptomic signatures, GxVAEs [18] jointly models bulk gene expression and molecular latent spaces, GexMolGen [19] aligns scRNA-seq foundation model-encoded gene expression representations with a scaffold-based molecular decoder, and MolGene-E [20] extends this direction toward single-cell chemical transcriptomics. However, MolGAN and GxVAEs lack single-cell evaluation, whereas GexMolGen and MolGene-E primarily integrate transcriptomic and molecular representations through two-stage alignment. Existing methods are also not explicitly designed to preserve cell type-specific transcriptional differences throughout molecular generation, have not systematically demonstrated generalization to unseen cell types, and have not been applied to personalized therapeutic discovery using patient-derived single-cell profiles. Here we introduce **GEM-GPT** (**G**ene **E**xpression-guided **M**olecular **G**enerative **P**re-trained **T**ransformer), which couples a pretrained scRNA-seq foundation model with a molecular GPT through biology-inspired deep fusion. GEM-GPT conditions autoregressive molecular generation on the desired transition from a disease-associated source cell state toward a matched healthy target state, and repeatedly integrates transcriptomic information throughout the molecular generator rather than compressing cellular context into a single fixed conditioning vector. We evaluate whether this design learns generalizable transcriptome-to-molecule relationships across bulk and single-cell-derived chemical perturbations, unseen cell lines, and CRISPR knock-out signatures. We further apply GEM-GPT to opioid use disorder (OUD), a heterogeneous chronic disorder for which available treatments remain limited [30–32] and patient- and cell type-specific transcriptional responses have been reported [33, 34]. Across these settings, GEM-GPT produces distinct molecular outputs across cellular contexts, generalizes to previously unseen cellular contexts and perturbation modalities, and identifies both novel candidates and FDA-approved drugs with OUD-relevant pharmacology. These capabilities establish a framework for personalized, cell type-resolved therapeutic design in systems pharmacology.

## 2 Results

### 2.1 Overview of GEM-GPT

GEM-GPT uses a GPT-based autoregressive architecture to generate SMILES strings conditioned on a desired transcriptional transition between a source cellular state and a target cellular state, corresponding in therapeutic applications to a disease-associated state and its healthy counterpart (Fig. 1a). During generation, a pretrained and frozen scRNA-seq foundation model, scGPT [23], encodes the paired diseased and healthy gene expression profiles into layer-specific transcriptomic representations. We use scGPT because its cell-level embeddings capture cellular identity and generalize across diverse single-cell datasets [23]. A scGPT-derived cell state embedding initializes molecular generation, while a deep fused attention mechanism couples the molecular hidden state with healthy- and disease-state representations at every layer (Fig. 1b), allowing the desired disease-to-healthy transcriptional shift to guide generation throughout the network. Full architectural details are provided in Methods.

**Fig. 1.**
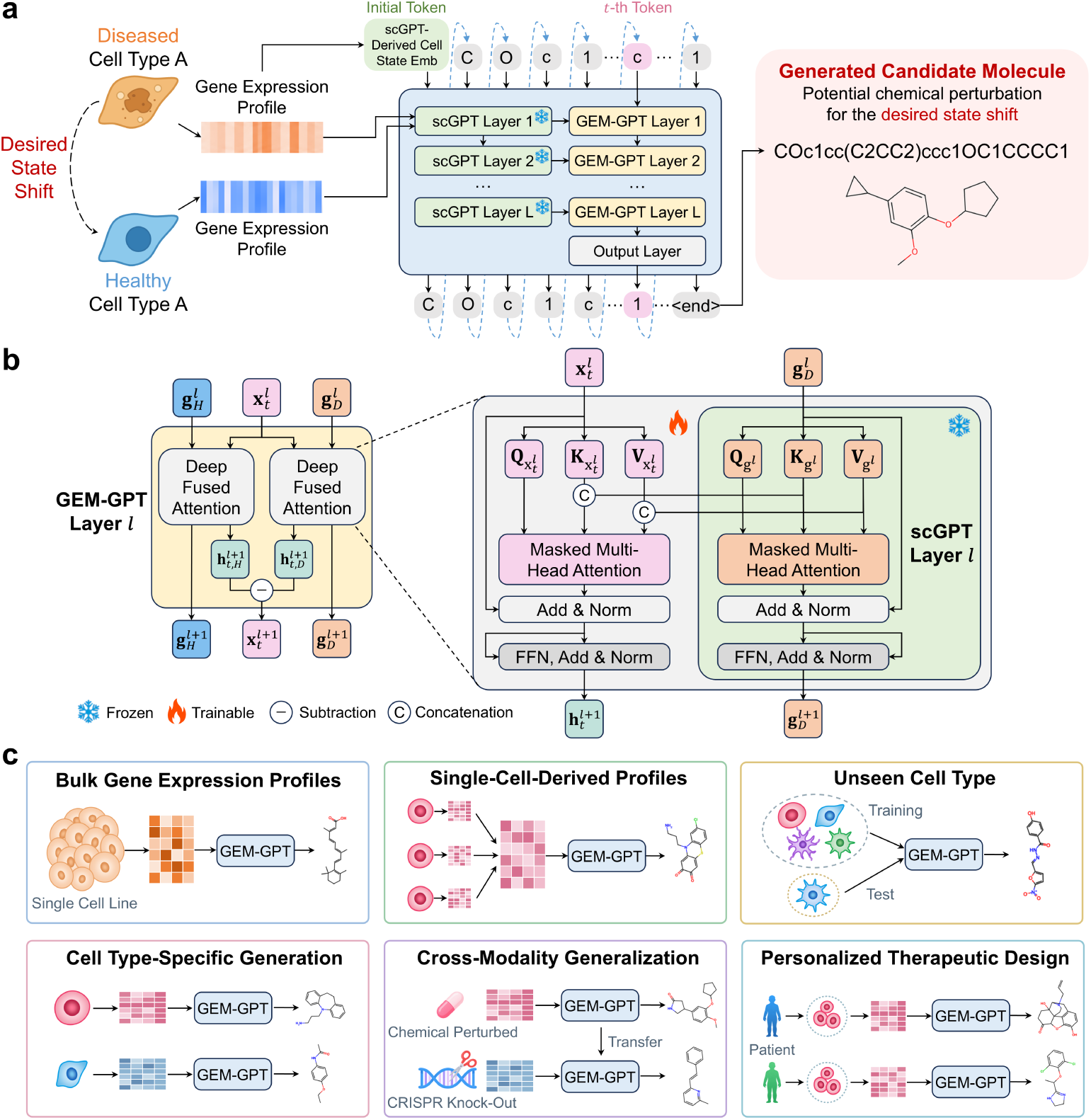
Overview of GEM-GPT. **a.** GEM-GPT autoregressively generates molecular SMILES strings by conditioning on the transcriptional transition from a diseased cell state toward its healthy counterpart. A pretrained scRNA-seq foundation model, scGPT, encodes the diseased and healthy gene expression profiles, while the scGPT-derived cell state embedding serves as the initial token embedding for cell type-specific molecular generation. **b.** Detailed architecture of a GEM-GPT block and the proposed deep fused attention mechanism. At each layer, the embedding of the *t*-th molecular token, **x***t*, is separately fused with the corresponding scGPT hidden representations of the healthy and diseased gene expression embeddings **g***_H_* and **g***_D_*. The difference between the resulting conditioned molecular representations is propagated to the next GEM-GPT layer, allowing the desired disease-to-healthy transcriptional shift to guide molecular generation throughout the network. The enlarged view illustrates the layer-level fusion between the trainable molecular attention module and the frozen scGPT attention module. **c.** Overview of the biological settings and capabilities evaluated for GEM-GPT, including molecular generation from bulk gene expression profiles and single-cell-derived profiles, generalization to unseen cellular contexts, cell type-specific molecular generation, cross-modality generalization from chemical perturbations to CRISPR knock-out signatures, and personalized therapeutic design using patient-derived transcriptomic profiles.

We evaluated GEM-GPT against state-of-the-art baselines [17–20] across four experiments, covering the biological contexts and capabilities summarized in Fig. 1c. We first tested hit-like molecule generation from bulk compound-induced profiles in seen and unseen cell lines. We then transferred the models to single-cell-derived chemical perturbation profiles to assess molecular generation. We next investigated whether GEM-GPT could generalize from chemical perturbations to CRISPR knock-out signatures. Finally, we applied GEM-GPT to patient-derived single-nucleus RNA-seq data from individuals with OUD to generate therapeutic candidates, evaluating its potential for personalized and cell type-specific therapeutic design.

### 2.2 GEM-GPT enables cell type-specific hit-like molecule generation from gene expression profiles across seen and unseen cell lines

A central objective of GEM-GPT is to move beyond the conventional one-drug-one-target paradigm by learning the relationship between chemical structures and systems-level transcriptional phenotypes. Rather than conditioning molecular generation on an isolated molecular target, GEM-GPT uses gene expression profiles to represent cellular responses to chemical perturbations. We therefore evaluated the model on L1000 compound-induced profiles under two benchmarks to isolate different axes of generalization: an in-distribution (ID) cell-line setting with a compound-based split, which tests unseen chemical perturbations in represented cell lines, and an out-of-distribution (OOD) cell-line setting in which test cell lines were excluded from training (Fig. 2a, c; Methods). The ID setting tests chemical generalization, whereas the OOD setting directly tests whether the learned transcriptome-to-molecule relationship transfers to cellular populations that were never observed during model development.

**Fig. 2.**
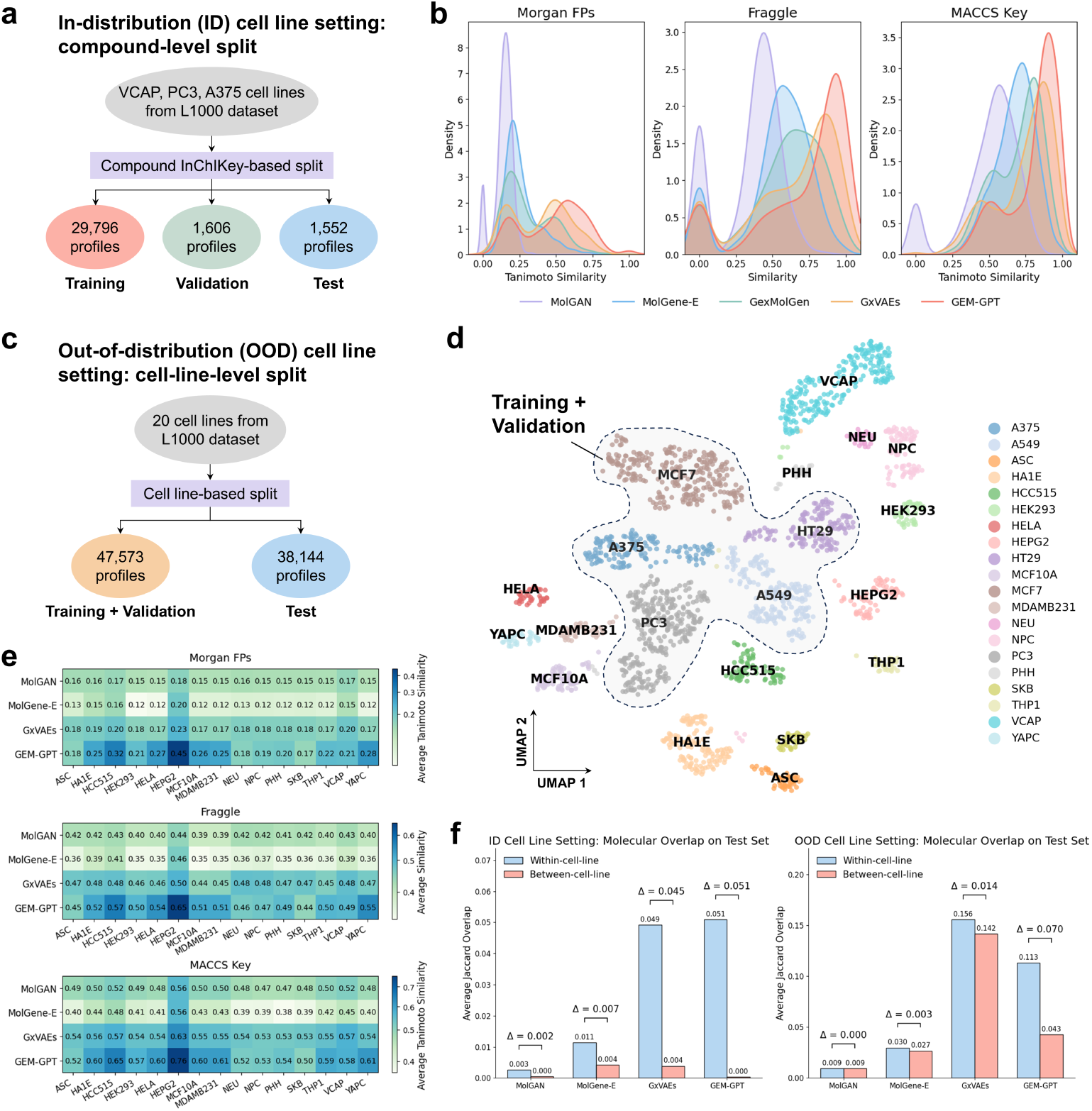
GEM-GPT generates cell type-specific hit-like molecules from bulk compound-induced gene expression profiles in both in-distribution and out-of-distribution cell-line settings. **a.** Schematic of the in-distribution (ID) cell-line setting using L1000 compound-induced gene expression profiles from 3 cell lines with a compound InChIKey-based split. **b.** Kernel density estimates of the similarities between generated molecules and their corresponding reference compounds in the ID setting. **c.** Schematic of the out-of-distribution (OOD) cell-line setting using L1000 compound-induced gene expression profiles from 20 cell lines with a cell-line-level split. **d.**UMAP visualization of control gene expression profiles from the 20 cell lines in the OOD setting. Dashed contours indicate the training and validation cell lines. **e.** Heatmaps of the average similarities between generated molecules and their corresponding reference compounds for each OOD test cell line and model. **f.** Mean Jaccard overlap between molecular sets generated within the same cell line and between different cell lines in the ID and OOD settings. Δ denotes the within-cell-line minus between-cell-line overlap. A larger Δ indicates a greater separation between within-cell-line and between-cell-line molecular overlaps, supporting stronger cell type-specific differentiation of the generated molecular sets.

In the ID setting, GEM-GPT achieved the highest validity and uniqueness among the evaluated models and the highest QED and lowest SA, while maintaining high novelty and internal diversity (Supplementary Fig. 1). Because these metrics do not directly measure whether the generated molecule corresponds to the input transcriptional perturbation, we followed previous transcriptomics-guided molecular generation studies [17–20] and treated structural similarity to the corresponding reference compound as the primary phenotype-to-chemistry evaluation. GEM-GPT produced molecules with consistently greater similarity to their corresponding reference compounds than the baselines across Morgan fingerprint-based Tanimoto, fragment-based Fraggle, and MACCS structural key-based Tanimoto similarity (Fig. 2b). The consistent improvement across diverse chemical structure representations supports a stronger correspondence between the input transcriptional perturbation and multiple structural characteristics of the generated molecule.

In the OOD setting, the UMAP visualization showed that test cell lines occupied transcriptional regions distinct from those represented during training and validation (Fig. 2d). Under this more challenging setting, the models exhibited distinct trade-offs across the conventional molecular generation and drug-likeness metrics (Supplementary Fig. 2). GEM-GPT maintained the highest validity among all methods, while no single model consistently outperformed the others across the remaining metrics. GEM-GPT nevertheless achieved the highest overall similarities to the corresponding reference compounds across the OOD test cell lines and all three structural representations (Fig. 2e). Thus, GEM-GPT transferred the learned transcriptome-to-molecule relationship to cellular contexts absent from model training. This property is particularly important for personalized design because clinically relevant cell types and patient states may be absent in available training datasets.

To assess cell type-specific molecular generation, we compared the mean Jaccard overlaps of molecular sets generated within the same cell line and between different cell lines in both benchmarks (Fig. 2f). A model that collapses toward generic molecules would show high overlap regardless of cell line, whereas a model that generates highly variable molecules without retaining cellular information would show uniformly low overlap. GEM-GPT showed the largest separation between within-cellline and between-cell-line overlap, with Δ = 0.051 in the ID setting and Δ = 0.070 in the OOD setting, indicating preserved within-cell-line consistency while differentiating chemical space across cell lines. In contrast, the high overlaps of GxVAEs in the OOD setting indicated weaker cell-line differentiation, whereas the near-zero overlaps of MolGAN and MolGene-E in both settings indicated highly diverse generated molecules regardless of cellular context.

To further investigate which architectural components support this performance, we performed an ablation study in the OOD setting (Supplementary Table 1). Removing deep fused attention while retaining the initial cell state embedding reduced performance, showing that input-level conditioning alone was insufficient. Removing all scGPT-related components produced a larger degradation, and removing only the initial cell state embedding caused the largest performance loss. Replacing the two parallel healthy- and disease-conditioned branches with a single concatenation-based fusion strategy also reduced performance. These results support the roles of early cell-state conditioning, pretrained single-cell representations, and deep fusion in GEM-GPT. Together, the L1000 experiments show that GEM-GPT learns a generalizable mapping between transcriptional changes and chemical structures while preserving molecular patterns associated with cellular context.

### 2.3 GEM-GPT extends hit-like molecule generation from bulk to single-cell compound-induced gene expression profiles

Following the bulk experiments, we investigated transcriptomics-guided molecular generation using perturbation profiles derived from single-cell measurements, which capture cell context-specific transcriptional responses [35, 36]. We evaluated GEM-GPT and baselines on sci-Plex3 [36], which contains profiles induced by 129 compounds. Due to the limited number of compounds, all models were initialized from L1000-trained models and continued training on sci-Plex3, whose cell lines are not among those used for L1000 training. Training on sci-Plex3 used a compound-based split with non-overlapping compounds across training, validation, and test sets (Methods).

GEM-GPT retained competitive conventional molecular generation properties after transfer to the sci-Plex3 dataset, ranking second in validity, uniqueness, IntDiv, and SA, while no model consistently outperformed the others across these metrics (Supplementary Fig. 3). More importantly, distributions of Morgan fingerprint-based, fragment-based Fraggle, and MACCS Tanimoto similarity were consistently shifted toward higher values than those of the baselines (Fig. 3a). Notably, GxVAEs achieved the highest uniqueness, IntDiv, and QED but the lowest similarity to the corresponding reference compounds across all three measures, underscoring that conventional generation metrics alone do not establish correspondence to the input transcriptional perturbation.

**Fig. 3.**
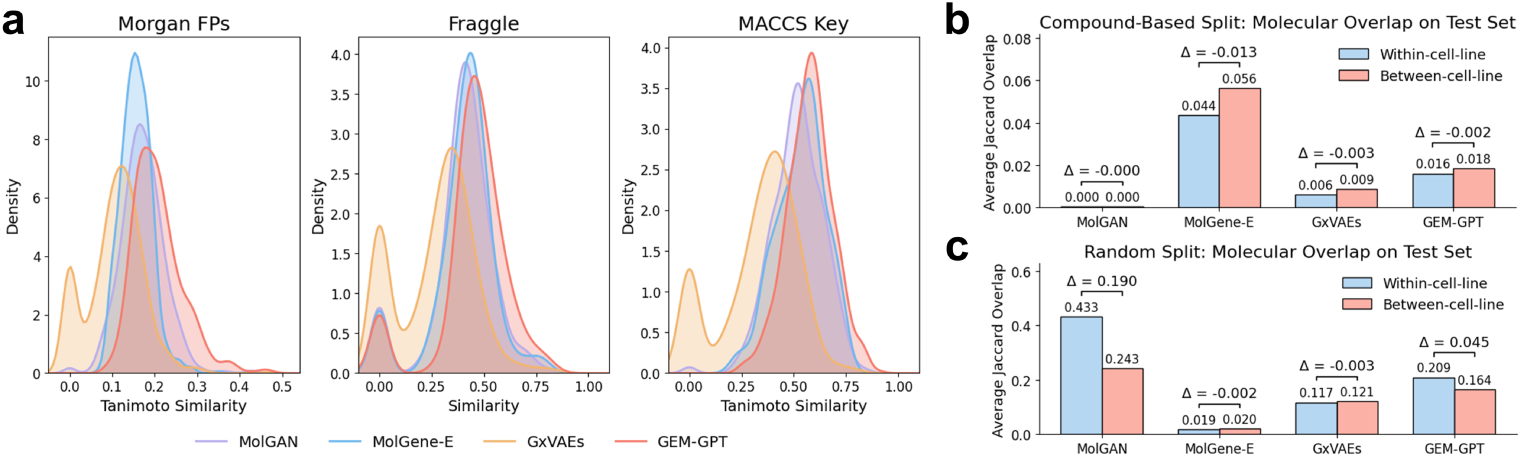
GEM-GPT extends hit-like molecule generation from bulk to single-cell compound-induced gene expression profiles. **a.** Kernel density estimates of the similarities between generated molecules and their corresponding reference compounds in sci-Plex3. **b-c.** Mean Jaccard overlap between molecular sets generated within the same cell line and between different cell lines under the compound-based split (**b**) and random split (**c**). Δ denotes the within-cell-line minus between-cell-line overlap.

We next asked whether the generated chemical space retained cell-line-specific information. Under the stringent compound-based split, Δ was non-positive for all models, and, except for MolGene-E, both within-cell-line and between-cell-line overlaps were below 0.02 (Fig. 3b). We reasoned that compound-level generalization under the sci-Plex3 compound-based split, combined with the limited chemical coverage of sci-Plex3, makes repeated cell-line-associated molecular patterns difficult to learn. In a diagnostic random-split experiment allowing repeated compound-associated perturbation information across training and test data, both GEM-GPT and MolGAN exhibited a positive Δ (Fig. 3c). Together, these results extend GEM-GPT from bulk profiles to single-cell-derived transcriptomics, while showing stronger structural correspondence to reference compounds and suggesting that cell-line-specific molecular patterns can emerge when sufficient repeated perturbation information is available.

### 2.4 GEM-GPT generates pharmacologically relevant molecules from CRISPR knock-out transcriptomic signatures

Since GEM-GPT is designed to translate transcriptional states into candidate molecules, we next investigated whether it could generate pharmacologically relevant structures from genetically perturbed cellular states. Models trained only on compound-induced L1000 data were evaluated, without additional training, on CRISPR knock-out signatures for 10 genes with known inhibitors in ExCAPE [37] (Methods). This setting tests whether the learned transcriptome-to-molecule mapping transfers from chemical to genetic perturbations.

Across the 10 genes, GEM-GPT generally generated molecules that were more similar to the corresponding known inhibitors than those produced by MolGAN, MolGene-E, GexMolGen, and GxVAEs (Fig. 4a). The trend was consistent across Morgan fingerprint-based Tanimoto, Fraggle, and MACCS Tanimoto similarity, indicating recovery of inhibitor-associated structural characteristics rather than features specific to one molecular representation.

**Fig. 4.**
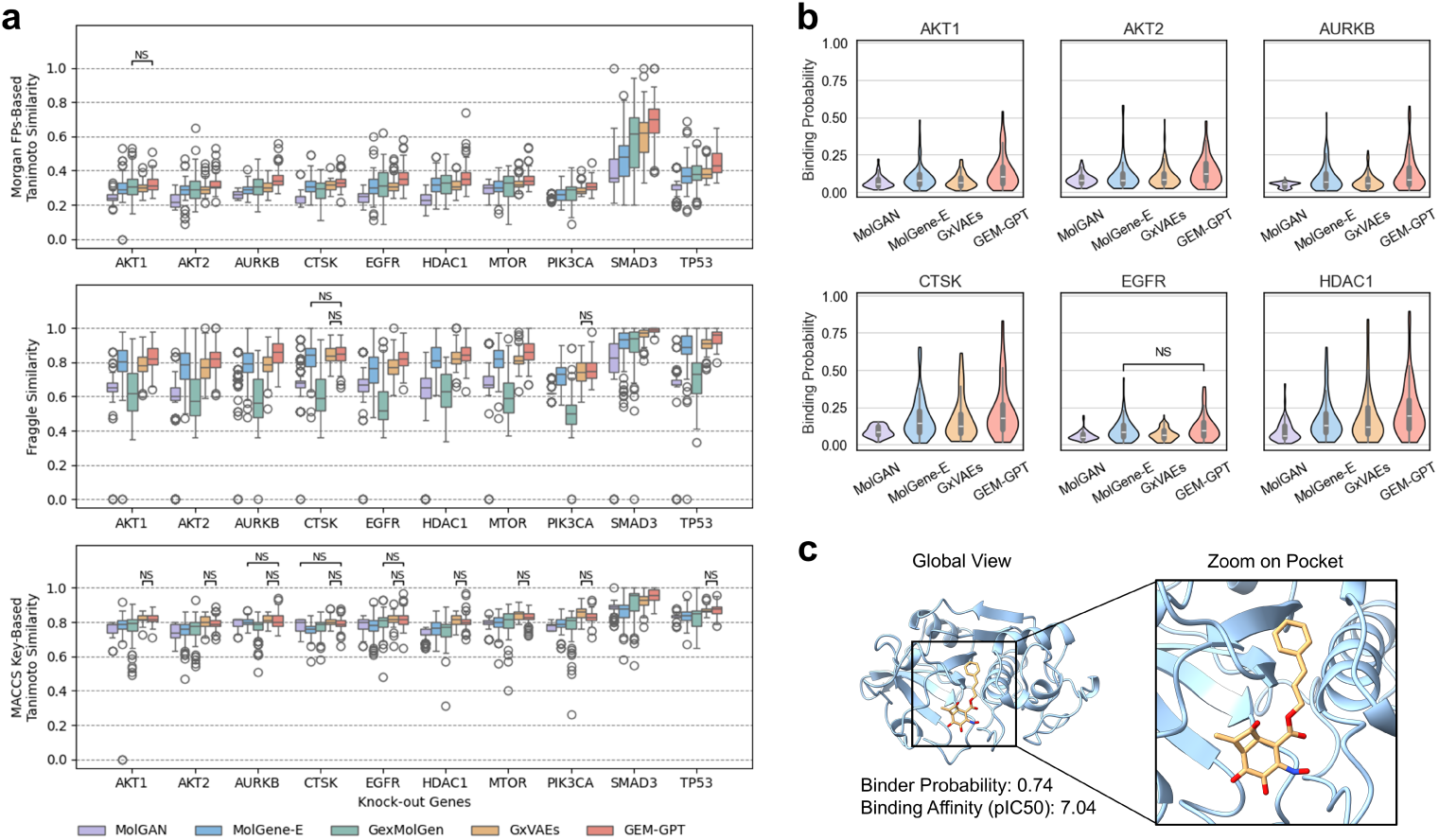
GEM-GPT generates potential gene inhibitors from bulk CRISPR knock-out transcriptomic signatures. **a.** Box plots of similarities between molecules generated from the knock-out signatures of 10 human genes and known inhibitors of their corresponding protein targets. GEM-GPT generally generated molecules with greater structural similarity to the corresponding known inhibitors than the baseline methods. NS indicates that the difference between GEM-GPT and the corresponding baseline was not statistically significant, with *P ≥* 0.05. **b.** Violin plots of Boltz-2-predicted binding probabilities for the subset of knocked-out genes whose corresponding protein targets are druggable enzymes with a single binding pocket. GEM-GPT generally generated molecules with higher predicted binding probabilities against the corresponding protein targets than the baseline methods. **c.** Representative example of a potential CTSK inhibitor generated by GEM-GPT. The generated molecule exhibits strong Boltz-2-predicted binding to CTSK.

Since structural similarity does not directly establish target interaction, we further evaluated generated molecules with Boltz-2 [38]. For the druggable enzymes among the 10 gene-associated targets, GEM-GPT generated molecules with higher mean predicted binding probabilities than each baseline across all evaluated targets, with most pairwise differences reaching statistical significance (Fig. 4b and Supplementary Table 2). A representative molecule generated from the CTSK knock-out signature achieved a predicted binder probability of 0.74 and a pIC50 of 7.04 for CTSK, where pIC50 is defined as *−* log_10_(IC50, [M]) (Fig. 4c).

These results are notable because GEM-GPT received neither target identities nor known inhibitor structures during generation and had never been trained on genetic perturbation data. Together, the agreement between inhibitor similarity and predicted binding highlights GEM-GPT’s cross-modality generalization from chemical to genetic perturbations and its potential to generate candidate modulators of corresponding protein targets for real-world drug discovery.

### 2.5 GEM-GPT generates personalized and cell type-specific therapeutic candidates for opioid use disorder using patient-derived single-nucleus transcriptomic data

To evaluate the utility of GEM-GPT in a clinically relevant setting, we established a computational pipeline for personalized and cell type-specific therapeutic design in OUD (Fig. 5a). For each patient and cell type, GEM-GPT was conditioned on the transition from the patient-specific OUD expression profile toward a cell type-matched healthy reference profile, with the goal of generating molecules that could potentially shift the disease-associated state toward a healthy phenotype. Candidate sets were first analyzed for patient and cell type specificity. We then used the Similarity Ensemble Approach (SEA) [39] as a post-generation analysis to identify potential protein target associations, followed by Boltz-2 [38] to prioritize selected candidates by predicted binding. Detailed preprocessing, filtering thresholds, and downstream prioritization criteria can be found in Methods.

**Fig. 5.**
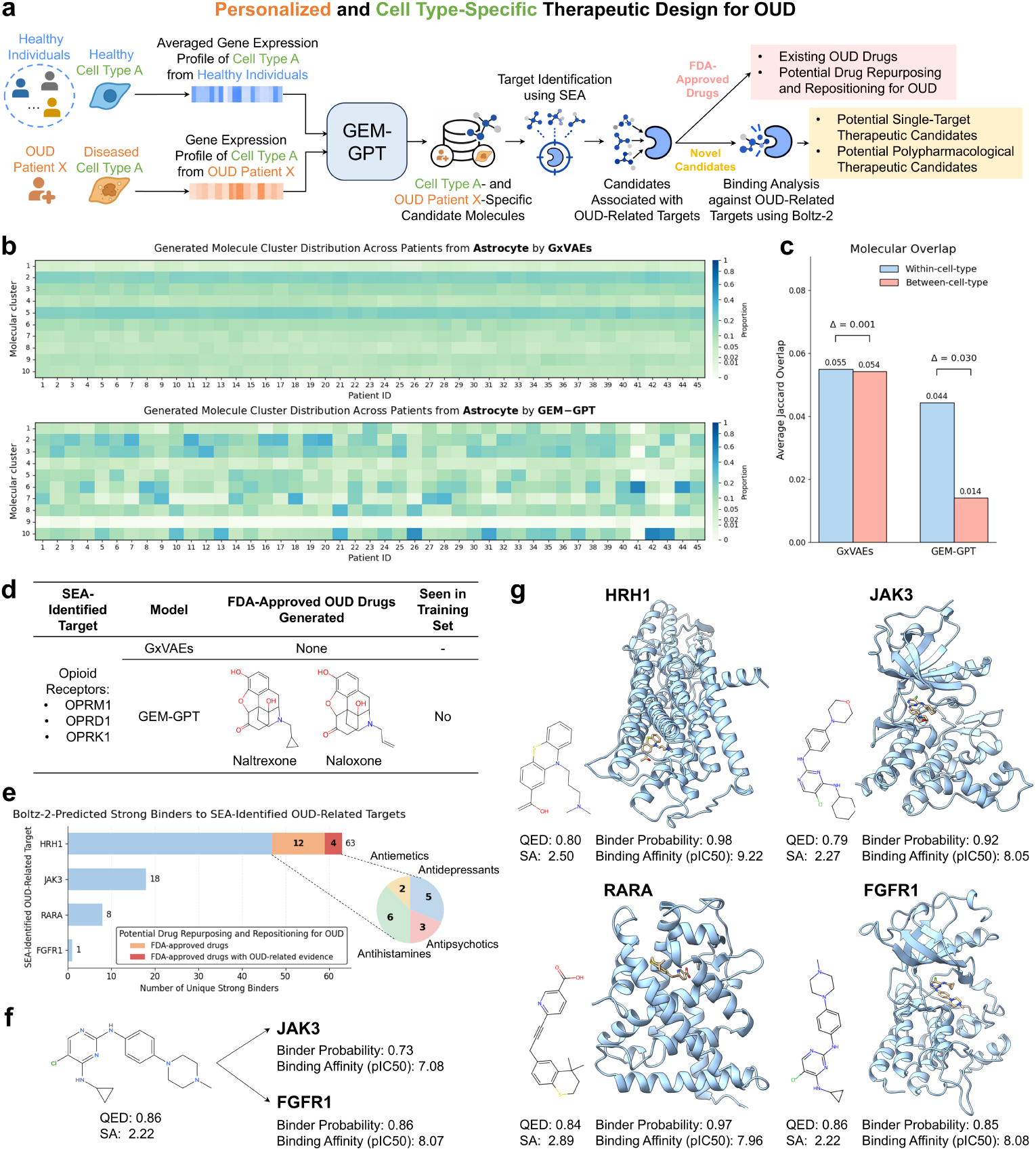
GEM-GPT enables personalized and cell type-specific therapeutic design for OUD. **a.** Pipeline for generating candidate molecules from paired cell type-specific gene expression profiles of OUD patients and healthy individuals, followed by post-generation target identification using Similarity Ensemble Approach (SEA) and binding analysis using Boltz-2. **b.** Molecular cluster distributions of candidates generated by GxVAEs and GEM-GPT across OUD patients using astrocyte-specific gene expression profiles. Differences in cluster distributions among patients reflect patient-specific molecular generation. **c.** Mean Jaccard overlap between molecular sets generated within the same cell type and across different cell types. Δ denotes the within-cell-type minus between-cell-type overlap. **d.** Among the generated molecules associated with SEA-identified opioid receptors (OPRM1, OPRD1, and OPRK1), GEM-GPT recovered 2 FDA-approved OUD-related drugs, neither of which appeared in the training set, whereas GxVAEs recovered none. **e.** Boltz-2 binding analysis of generated molecules associated with prioritized SEA-identified OUD-related targets (HRH1, JAK3, RARA, and FGFR1) whose corresponding genes are upregulated in our investigated cell types. Among the GEM-GPT-generated strong binders (binder probability *>* 0.8, pIC50 *>* 7) for HRH1, 16 are FDA-approved drugs and therefore represent potential drug repurposing or repositioning candidates for OUD; 4 of these have reported evidence for 1tr1eating OUD-related symptoms. **f.** Representative GEM-GPT-generated molecule predicted by Boltz-2 to bind both JAK3 and FGFR1, illustrating a potential polypharmacological therapeutic candidate. **g.** Representative GEM-GPT-generated strong binders for HRH1, JAK3, RARA, and FGFR1, with Boltz-2-predicted binding complexes and properties.

We applied this pipeline to single-nucleus RNA-seq profiles from postmortem ventral midbrain samples of individuals with OUD and opioid-free controls [40], focusing on four glial cell types: astrocytes, microglia, oligodendrocytes (ODC), and oligodendrocyte precursor cells (OPC). These populations were consistently represented across patients and showed substantial OUD-associated transcriptional alterations in the source dataset [40]. None was represented in the L1000 cell-line training set, providing an additional evaluation in previously unseen cellular populations. For each cell type, the opioid-free control reference was constructed from opioid-free controls and paired with each patient’s OUD profile. GxVAEs, the strongest overall baseline in our preceding experiments, was evaluated using expression signatures derived from the same inputs and generation protocol.

We first assessed general properties of the generated molecules (Supplementary Fig. 4). GEM-GPT achieved slightly lower validity than GxVAEs, but outperformed GxVAEs in uniqueness, novelty, internal diversity, and QED, and achieved a lower SA score. Thus, GEM-GPT generated a diverse and drug-like candidate space with favorable predicted synthetic accessibility while maintaining high validity.

To assess inter-patient specificity, generated molecules were represented using Morgan fingerprints and grouped into molecular clusters. If patient-specific transcriptomic information is propagated through generation, diverse patients within the same cell type should occupy different proportions of the chemical clusters. GEM-GPT showed distinct cluster distributions across patients within astrocytes and the other investigated cell types (Fig. 5b and Supplementary Fig. 5), whereas GxVAEs produced substantially more similar distributions. These patterns indicate that GEM-GPT translates inter-patient differences into differentiated candidate chemical spaces rather than generating a uniform population-level output.

We next evaluated cell type specificity by comparing molecular overlap within and across cell types for the same patient. GxVAEs showed nearly the same within-cell-type and between-cell-type overlaps, resulting in Δ = 0.001 (Fig. 5c). GEM-GPT instead got Δ = 0.030. The within-cell-type overlap of GEM-GPT remained modest (0.044), consistent with chemical diversity, but the substantially lower cross-cell-type overlap (0.014) demonstrates that the generated molecular space is more strongly structured by cell identity. Together with the patient-level cluster analysis, these results support both personalized and cell type-resolved generation in patient-derived data.

SEA-based target analysis provided complementary pharmacological evidence. For opioid receptors, we focused on OPRM1, OPRD1, and OPRK1, which are central to opioid pharmacology and are all readily detectable among the four glial cell types we investigated [40]. Notably, all three were found among the SEA-identified targets associated with GEM-GPT-generated molecules. Among these molecules, GEM-GPT recovered naltrexone and naloxone (Fig. 5d). Naltrexone is FDA-approved for OUD treatment, and naloxone is an FDA-approved opioid antagonist used for emergency overdose reversal. Neither molecule appeared in the GEM-GPT training set, and GxVAEs recovered no FDA-approved OUD-associated drug under the same analysis. The recovery of these clinically established compounds provides retrospective pharmacological evidence that GEM-GPT can generate molecules consistent with established opioid pharmacology from disease-to-reference transcriptomic conditioning, without being provided receptor identities or OUD drug structures during generation.

Beyond the classical opioid receptors, we prioritized four SEA-identified targets, HRH1, JAK3, RARA, and FGFR1, whose genes are upregulated in the analyzed glial cell types [40] and have reported relationships to OUD or opioid-associated biology [41–46]. After stringent SEA filtering and Boltz-2 evaluation, GEM-GPT generated strong predicted binders for all four targets (Fig. 5e,g). HRH1 provided a particularly notable repurposing example: 16 of 63 strong predicted HRH1 binders were FDA-approved drugs spanning antihistamine, antidepressant, antipsychotic, and antiemetic classes (Supplementary Table 3). Four of these, amitriptyline, imipramine, doxepin, and prochlorperazine, have reported evidence related to OUD or opioid withdrawal-associated symptoms [47–50]. Although these observations do not establish therapeutic efficacy, they identify clinically characterized compounds for targeted follow-up and illustrate how transcriptome-guided generation can complement *de novo* design with repurposing and repositioning hypotheses.

Finally, we investigated whether GEM-GPT could generate novel candidates with potential polypharmacological activity. Among candidates associated with the four prioritized targets, four GEM-GPT-generated molecules were predicted by Boltz-2 to bind more than one target under the specified multi-target criteria (Supplementary Table 4). A representative molecule was predicted to bind both JAK3 and FGFR1 with favorable binder probabilities and affinities (Fig. 5f). Such multi-target candidates may be particularly relevant to systems pharmacology, where coordinated modulation of multiple disease-associated pathways could be advantageous.

Together, the OUD case study shows that GEM-GPT can generate patient-specific and cell type-specific chemical spaces for novel cell types, recover clinically relevant drugs absent from training, and nominate both single-target and polypharmacological candidates for downstream experimental validation.

## 3 Discussion

In this study, we developed GEM-GPT, a transcriptomics-guided cell type-resolved molecular generation framework that integrates a pretrained single-cell foundation model with an autoregressive molecular language model through deep fused attention. GEM-GPT was designed to support systems pharmacology by generating cell type-specific candidate molecules from desired transitions between disease-associated and healthy cellular states rather than conditioning generation on a predefined molecular target. Across bulk and single-cell chemical perturbation profiles, CRISPR knock-out signatures, unseen cell lines, and patient-derived OUD profiles, the model generated molecules that preserved information associated with the input transcriptional context and generalized beyond the cellular and perturbational settings used for training.

A central contribution of GEM-GPT is its unified modeling of transcriptomic and molecular representations during autoregressive generation. Existing transcriptomics-guided approaches either condition generation on bulk expression representations or align transcriptomic and molecular latent spaces through separate objectives. In contrast, GEM-GPT incorporates transcriptomic information throughout the molecular generator, allowing the desired disease-to-healthy transition to influence successive molecular hidden states. The ablation results support both the initial cell state embedding and repeated deep fusion as contributors to performance.

In the compound-induced experiments, molecules generated by GEM-GPT showed greater similarity to reference compounds in both seen and unseen cell lines and a larger separation between within-cell-line and between-cell-line molecular overlap, supporting both transferability and cellular specificity. The sci-Plex3 experiments extended the structural correspondence to single-cell perturbation data, while also showing that limited chemical coverage can constrain cell-line-specific patterns. The CRISPR experiment further demonstrated cross-modality generalization, with molecules generated from knock-out signatures showing greater similarity to known inhibitors and higher predicted binding probabilities for corresponding druggable targets, despite GEM-GPT being trained only on compound-induced profiles. Finally, the OUD case study extended GEM-GPT to patient-derived disease data, where distinct candidate distributions across patients and cell types, the recovery of FDA-approved drugs relevant to OUD care absent from training, and the identification of repurposing and multi-target candidates demonstrated the model’s ability to generate clinically relevant molecular candidates in a patient- and cell-type-specific manner.

Several limitations should be considered. First, GEM-GPT inherits limitations of the available perturbational data. L1000 measures a restricted landmark-gene set and is dominated by immortalized cell lines, whereas current single-cell chemical perturbation datasets have limited compound coverage. A pretrained single-cell encoder can facilitate transfer but cannot eliminate the domain shift to primary patient-derived tissues. Second, our evaluations establish structural correspondence, molecular-set differentiation, and predicted target binding, but they do not demonstrate reversal of the intended disease phenotype. Experimental perturbation assays are required to test transcriptomic reversal, target engagement, toxicity, and phenotypic activity. Third, the current generation objective does not directly optimize selectivity, safety, metabolic stability, tissue exposure, or other development properties, which are particularly important for central nervous system indications such as OUD.

Future work should combine transcriptomic conditioning with larger cell type-resolved perturbation datasets, additional biological modalities, multi-objective optimization, uncertainty estimation, and iterative experimental feedback. Patient-derived cells, organoids, and in vivo models will be important for determining whether prioritized molecules produce reproducible therapeutic effects. Overall, GEM-GPT connects single-cell representations of disease state with transcriptomics-guided molecular generation and offers a path from target-centric design toward personalized, cell type-resolved, systems-oriented therapeutic discovery in complex diseases.

## 4 Methods

### 4.1 Architecture of GEM-GPT

GEM-GPT is a transcriptomics-conditioned autoregressive molecular generation framework designed to generate candidate compounds associated with a desired transition between a source cellular state and a target cellular state. In therapeutic applications, these states correspond to a disease-associated source state and a healthy target state. Given healthy and diseased gene expression profiles **G***^H^* and **G***^D^*, GEM-GPT generates a molecular SMILES sequence **s** = (*s*_1_*, s*_2_*, . . . , s_T_*). The conditional generation probability is factorized as

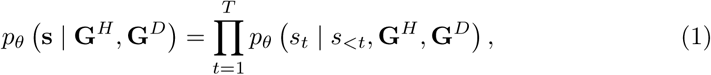

where *s_<t_*denotes the molecular tokens generated before position *t*, and *θ* denotes the trainable parameters of the molecular generator.

As shown in Fig. 1, GEM-GPT consists of a pretrained scRNA-seq foundation model, scGPT [23], and a GPT-based molecular generator. The healthy and diseased gene expression profiles are independently encoded by scGPT to obtain transcriptomic representations. The molecular generator then autoregressively predicts SMILES tokens while being conditioned on the desired transition between the two cellular states. The cell state embedding derived from scGPT is used as the initial embedding for molecular generation, allowing the generated molecular sequence to be conditioned on cellular state from the beginning of the autoregressive process.

The parameters of scGPT are frozen throughout model training to preserve the biological representations learned from large-scale single-cell transcriptomic data. The parameters of the molecular generator and the newly introduced fusion modules are trainable. Unlike two-stage approaches [19, 20] that independently encode transcriptomic and molecular modalities before aligning their latent representations, GEM-GPT directly integrates transcriptomic information into each layer of the molecular generator. This unified architecture allows transcriptomic and molecular representations to interact repeatedly throughout SMILES generation.

#### 4.1.1 Encoding gene expression profiles

Healthy and diseased gene expression profiles, **G***^H^* and **G***^D^*, are encoded separately using the pretrained scGPT model (`whole-human` version). For each profile, the input consists of gene identity tokens and their corresponding gene expression values. Genes are ordered according to the scGPT vocabulary, and genes absent from the pretrained vocabulary are excluded before model input.

Rather than using only the final scGPT representation, GEM-GPT retains the layer-specific hidden representations from every scGPT layer. The hidden gene expression representation **g** from each scGPT layer is supplied to the corresponding layer of the molecular generator through the deep fused attention mechanism. This design enables molecular generation to incorporate transcriptomic information represented at different levels of abstraction.

We also extract a cell state representation from scGPT by appending a <`cls`> token to the beginning of the input tokens. Following [23], the final-layer embedding corresponding to the <`cls`> token is extracted as the cell state representation. The scGPT-derived cell state embedding is used to initialize the molecular generation sequence, providing explicit cellular identity information, while the layer-specific scGPT representations provide continuous information about the healthy and diseased transcriptional states.

#### 4.1.2 Deep fused attention

GEM-GPT introduces a deep fused attention mechanism that integrates transcriptomic information into every layer of autoregressive molecular generation (Fig. 1b). At generation step *t* and molecular layer *l*, let **x***_t_^l^* denote the molecular hidden representation at the current SMILES position, and let **g***_H_^l^* and **g***_D_^l^* denote the layer-*l* scGPT representations of the healthy and diseased cellular states, respectively. The same molecular representation is processed by two parallel deep fused attention branches, one conditioned on **g***_H_^l^* and the other conditioned on **g***_D_^l^*.

The enlarged panel in Fig. 1b shows one of these two branches using a layer-*l* scGPT representation **g***^l^*. At autoregressive step *t*, the molecular branch processes the SMILES prefix available up to the current position. Let **X**_≤*t*_*^l^* = [ **x**_1_*^l^ ,* **x**_2_*^l^ , . . . ,* **x***_t_^l^*] denote the molecular hidden states available at layer *l*. The query representation for the current position is projected from **x***_t_^l^*, whereas the molecular key and value representations are projected from the available molecular prefix:

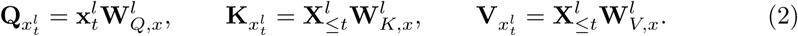

In parallel, the corresponding scGPT representation is projected into query, key, and value representations:

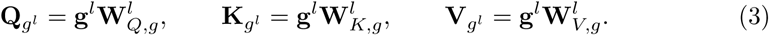

The scGPT query, key, and value representations are used by the frozen *l*-th scGPT layer to update the gene expression representation:

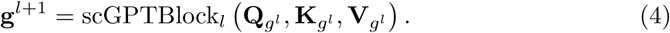

For molecular generation, the scGPT-derived keys and values are concatenated with the molecular keys and values along the sequence dimension:

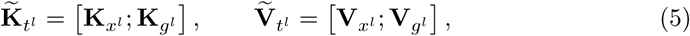

where [*·*; *·*] denotes concatenation. The molecular query remains derived exclusively from the current molecular representation. The fused attention output is calculated as

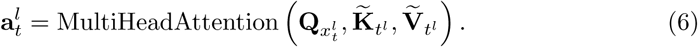

Masked multi-head attention is used in the molecular branch to preserve autoregressive generation, ensuring that the representation at position *t* does not access future molecular tokens. The fused attention output is subsequently processed by residual connections, layer normalization, and a feed-forward network:

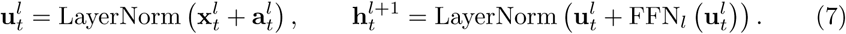

This deep fused attention operation is applied separately to the healthy and diseased gene expression representations to get healthy-conditioned molecular representation **h***_t,H_* and disease-conditioned molecular representation **h***_t,D_*:

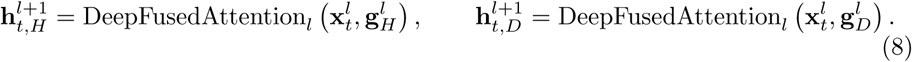

The molecular representation passed to the next GEM-GPT layer is obtained by subtracting the disease-conditioned representation from the healthy-conditioned representation:

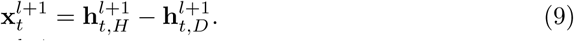

The resulting **x***_t_^l^*^+1^, **g***_H_^l^*^+1^, and **g***_D_^l^*^+1^ are supplied to the next GEM-GPT layer, where the same dual-branch fusion procedure is repeated. After the final layer, the molecular representation at position *t* is projected onto the SMILES vocabulary to predict the *t*-th token:

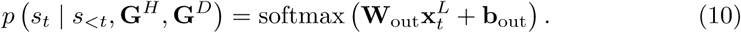

Through this layer-wise procedure, the representation used to generate each SMILES token is repeatedly conditioned on both healthy and diseased transcriptomic states. The subtraction between the two conditioned molecular representations explicitly encodes the desired disease-to-healthy transition at every molecular layer. The scGPT blocks remain frozen, whereas the molecular attention, feed-forward, and output modules are trainable.

#### 4.1.3 Autoregressive molecular generation and model training

Molecules are represented as SMILES strings and tokenized using a character-level vocabulary. Each sequence is surrounded by beginning-of-sequence and end-of-sequence tokens and padded to a maximum sequence length of 80 tokens.

GEM-GPT is formulated to generate molecules for a desired transition from a diseased state **G***^D^* toward a healthy state **G***^H^*. During training on chemical perturbation data, this formulation is instantiated by treating the matched control profile as the source state and the compound-perturbed profile as the target state, with the model learning to generate the compound associated with this transition. The model is trained to predict each SMILES token conditioned on the preceding molecular tokens and the paired transcriptional states using autoregressive negative log-likelihood:

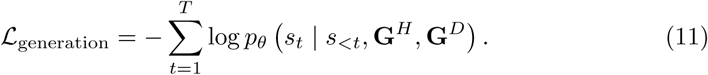

Only the molecular generator and deep fused attention modules are updated during optimization; pretrained scGPT parameters remain frozen. The model is optimized using AdamW [51], and the checkpoint with the lowest validation loss is selected for molecular generation. During inference, molecules are generated autoregressively using beam search until an end-of-sequence token or the maximum sequence length is reached. Generated SMILES strings are processed using RDKit [52]; strings that cannot be parsed or sanitized are treated as invalid, and valid molecules are converted to canonical SMILES for downstream analyses.

### 4.2 Experiments

The following subsections describe the experimental settings used to evaluate GEM-GPT in detail. All experiments were conducted on an NVIDIA L40S GPU with 48 GB of memory. The hyperparameter configurations used across the different experimental settings are summarized in Supplementary Table 5.

#### 4.2.1 Hit-like molecule generation from compound-induced transcriptomic profiles

Compound-induced transcriptomic data were obtained from the CMap LINCS 2020 L1000 dataset [53] and the sci-Plex3 dataset [36]. For the L1000 experiments, we used the 978 landmark genes directly measured by the platform. Gene expression profiles used as direct model inputs were normalized by library size and transformed using log(1 + x). Compound structures were standardized and converted to canonical SMILES using RDKit. The specific dataset construction, input representation, and data splitting strategies used in the different experimental settings are described below.

##### Bulk in-distribution cell-line setting

For the ID cell-line setting, we used the L1000-based dataset curated by [19], containing Level 3 compound-induced profiles from VCAP, PC3, and A375. GEM-GPT was compared with MolGAN, GxVAEs, GexMolGen, and MolGene-E. Because MolGAN, GxVAEs, and MolGene-E use differential expression signatures as input, we followed GxVAEs and computed each signature as log(*x*_cp_*/x*_ctl_), where *x*_cp_ and *x*_ctl_ denote the matched compound-perturbed and control expression values, respectively. The resulting differential expression signatures were used as inputs to these baseline models. Thus, all models used matched underlying perturbations. Samples were divided into training, validation, and test sets using a compound InChIKey-based split, with test compounds excluded from the training and validation sets. This setting was designed specifically to evaluate generalization to unseen compounds in represented cellular contexts. All models were evaluated under the same split and generated the same number of candidate molecules for each test profile.

##### Bulk out-of-distribution cell-line setting

For the OOD cell-line setting, we constructed a 20-cell-line dataset from CMap LINCS 2020 [53]. Because MolGAN, GxVAEs, and MolGene-E use differential expression signatures as input, whereas GEM-GPT requires paired control and compound-perturbed gene expression profiles, we constructed matched datasets from the Level 5 and Level 3 data, respectively. Level 5 differential expression signatures were used for the baseline models, whereas the corresponding Level 3 control and perturbed profiles were used for GEM-GPT, with a one-to-one correspondence maintained between the two datasets to ensure that all models were trained and evaluated on the same chemical perturbations. To construct the Level 5 dataset, we first retained chemical perturbations with doses of 5 or 10 *µ*M and a perturbation time of 24 h, following the setting of [17]. Using the replicate information recorded in distil ids, we excluded Level 5 signatures derived from fewer than two samples and then selected the 20 cell lines with the largest numbers of remaining signatures. Perturbations were further excluded if the associated molecule was invalid, could not be converted to canonical SMILES, or had a canonical SMILES length *>* 80 characters. We additionally removed Level 5 signatures whose experimental plates could not be identified in the Level 3 data. To construct the corresponding Level 3 dataset, the sample identifiers in distil ids for each retained Level 5 signature were matched to sample id entries in the Level 3 chemical perturbed data. For each matched perturbed sample, corresponding Level 3 control samples were identified according to plate ID, cell line, and perturbation time. Level 3 perturbed samples associated with the same Level 5 signature were averaged to obtain a single perturbed profile, while the corresponding control samples were averaged separately to obtain a matched control profile. The resulting Level 3 dataset therefore contained the same amount of data as the Level 5 dataset, with each Level 5 differential expression signature corresponding to one paired Level 3 control and perturbed profile. The final matched dataset contained data from 89,615 entries in total. Among the resulting 20 cell lines, the five cell lines with the largest numbers of available profiles, A375, A549, HT29, MCF7, and PC3, were used for training and validation, whereas the remaining 15 cell lines, ASC, HA1E, HCC515, HEK293, HELA, HEPG2, MCF10A, MDAMB231, NEU, NPC, PHH, SKB, THP1, VCAP, and YAPC, were excluded from model training and used as the held-out test set. The held-out factor in this benchmark was cell-line identity rather than compound identity, as chemical generalization was evaluated separately in the ID setting. The number of profiles for each cell line and the corresponding data split are shown in Supplementary Fig. 6. GEM-GPT was compared with MolGAN, GxVAEs, and MolGene-E using the matched datasets and the same generation scale. To evaluate cell-line-specific molecular generation, molecular sets generated within the same cell line were compared with those generated across different cell lines. Generation instances were randomly divided by input profile, ensuring that all molecules generated from the same profile remained together in each division, and the procedure was repeated five times. Mean Jaccard overlap was calculated within and between cell lines, and Δ was defined as the within-cell-line overlap minus the between-cell-line overlap.

##### Single-cell setting

Single-cell compound-induced gene expression profiles were obtained from sci-Plex3 [36]. Genes with zero counts across all cells were removed before denoising. Negative binomial and zero-inflated negative binomial (ZINB) distributions were compared for each gene, and the ZINB model was selected because the majority of genes favored zero inflation. The count matrix was then denoised and imputed using a Deep Count Autoencoder [54] with a ZINB output distribution, with genes dropped during denoising restored by zero-filling. Chemical-perturbed cells were restricted to those with more than 1,000 counts and valid compound annotations. For each cell line, timepoint, and plate, control cells were averaged to construct a matched control profile, and perturbed cells without a corresponding control group were excluded. Chemical-perturbed cells were then pseudo-bulked by averaging cells with the same cell line, timepoint, plate, compound, and dose, and paired with the corresponding control profiles. UMAP visualization of the processed profiles showed that cell type-specific transcriptional differences were retained after processing, with profiles from different cell types forming distinct clusters (Supplementary Fig. 7). The final analysis included 3,382 profiles induced by 129 compounds across MCF7, A549, and K562 cells. Because the number of unique compounds was insufficient for training the generative models from scratch, each model was initialized from the corresponding model trained on the dataset in the bulk ID cell-line setting and then continued training on sci-Plex3. None of the three sci-Plex3 cell lines overlapped with the VCAP, PC3, and A375 cell lines used for L1000 training. This transfer setting was designed primarily to evaluate adaptation from bulk transcriptomics to profiles derived from single-cell transcriptomics in previously unseen cell lines. The primary split was a 6:2:2 compound-based split with non-overlapping compounds across training, validation, and test sets. MolGAN, MolGene-E, GxVAEs, and GEM-GPT were trained and evaluated using the same split and transfer-learning strategy. For the diagnostic random-split analysis, profiles rather than compounds were randomly assigned to training, validation, and test sets, allowing profiles associated with the same compound to appear across splits. All models were retrained using the same random split, and within-cell-line and between-cell-line Jaccard overlaps were calculated using the same molecular-set procedure as in the L1000 experiments.

#### 4.2.2 Molecule generation from CRISPR knock-out transcriptomic signatures

GEM-GPT and the baseline models were trained on compound-induced L1000 profiles curated by [19] and evaluated on CRISPR knock-out signatures without additional training on gene knock-out data. This benchmark was designed to evaluate cross-modality transfer from chemical to genetic perturbations. Ten human genes with known inhibitors in ExCAPE [37] were selected: AKT1, AKT2, AURKB, CTSK, EGFR, HDAC1, MTOR, PIK3CA, SMAD3, and TP53. For each gene, 100 CRISPR knock-out profiles and 100 corresponding control profiles were used. Profiles were processed using the same 978 landmark genes, gene ordering, normalization procedure, and scGPT vocabulary mapping as the compound-induced experiment. An equal number of candidate molecules was generated for each gene, profile, and model.

Generated molecules were compared with known inhibitors using Morgan fingerprint-based Tanimoto similarity, Fraggle similarity, and MACCS key-based Tanimoto similarity. Protein-ligand binding was further evaluated using Boltz-2 [38] for the subset of knocked-out genes whose protein products were druggable enzymes with a single well-defined binding pocket: AKT1, AKT2, AURKB, CTSK, EGFR, and HDAC1. Each protein sequence and corresponding generated molecule were provided to Boltz-2 for binding probability prediction. Pairwise comparisons of predicted binding probabilities between GEM-GPT and each baseline were assessed using one-sided Mann-Whitney U tests, with P *<* 0.05 considered statistically significant.

#### 4.2.3 Personalized and cell type-specific therapeutic design for OUD

For the OUD application, we separately trained GEM-GPT on our curated L1000 compound-induced transcriptomic dataset covering 20 cell lines after excluding predefined OUD-related drugs from the training data. This model was trained using the same overall L1000 preprocessing and training framework described above and was subsequently applied to the OUD dataset without training or fine-tuning on any OUD patient-derived transcriptomic profiles. We applied GEM-GPT to single-nucleus transcriptomic profiles from 45 patients with OUD and 50 opioid-free controls [40]. The analysis focused on astrocytes, microglia, oligodendrocytes (ODC), and oligodendrocyte precursor cells (OPC), which were consistently represented across the patient samples and exhibited OUD-associated transcriptional alterations [40]. None of these brain cell types was included in the L1000 cell-line training set.

Expression values were normalized by total transcript count and transformed using log(1 + *x*). Gene identifiers were mapped to the scGPT vocabulary, and genes unavailable in the pretrained vocabulary were excluded. For each OUD patient and each cell type, all corresponding single-nucleus RNA-seq profiles were averaged to obtain one patient- and cell type-specific aggregated expression profile. For each cell type, single-nucleus RNA-seq profiles from all 50 opioid-free controls were averaged to construct a cell type-specific reference profile representing the desired healthy state. Despite aggregation, the resulting profiles retained clear cell type-specific transcriptional differences, as evidenced by the separation of the four cell types into distinct clusters in the UMAP visualization (Supplementary Fig. 8). Each model input therefore consisted of a patient-specific OUD profile paired with the corresponding cell type-specific healthy reference profile. GEM-GPT generated up to 1,000 molecular candidates for each pair of profiles. The same underlying patient-control profile pairs were used for GxVAEs, which received the corresponding differential-expression signatures according to its native input formulation and used the same generation scale.

Patient-specific molecular distributions were evaluated after converting valid canonical SMILES to Morgan fingerprints (radius=2, 2048 bits) and clustering the filtered molecules into 10 K-means clusters. For each patient and cell type, the fraction of generated molecules assigned to each cluster was calculated. Cell type-specific generation was evaluated by comparing Jaccard overlap between independently generated molecular sets within the same cell type and across different cell types in the same patient; random splitting was repeated five times.

For downstream SEA-based target search, only molecules with valid canonical SMILES were retained, and molecules with QED below 0.7 were excluded. SEA [39] was used after molecular generation to identify potential protein target associations from the chemical structures of the generated molecules. We first examined molecules associated with the classical opioid receptors OPRM1, OPRD1, and OPRK1. We then prioritized HRH1, JAK3, RARA, and FGFR1 based on their identification by SEA, expression in the investigated cell types, and reported relationships to OUD or opioid-associated biology [41–46]. For these four targets, molecule-target pairs were retained when the SEA *P* value was *<* 1 *×* 10*^−^*^10^ and the maximum Tanimoto coefficient to known target ligands was *>* 0.5. Retained pairs were evaluated with Boltz-2. Strong predicted binders were defined as molecules with binder probability *>* 0.8 and predicted pIC50 *>* 7. For the polypharmacology analysis, a molecule-target interaction was retained when binder probability was *>* 0.7 and predicted pIC50 was *>* 6; molecules satisfying these criteria for more than one prioritized target were classified as multi-target candidates.

#### 4.2.4 Molecular evaluation metrics

Conventional molecular generation metrics included validity, uniqueness, novelty, and internal diversity (IntDiv). Drug-likeness was assessed using quantitative estimate of drug-likeness (QED) and synthetic accessibility (SA) score. Transcriptomics-guided structural correspondence was evaluated using Morgan fingerprint-based Tanimoto similarity, Fraggle similarity, and MACCS key-based Tanimoto similarity to the corresponding reference compounds or known inhibitors. Molecular-set overlap was quantified by Jaccard similarity after canonicalization so that repeated representations of the same molecule were treated consistently.

## Data availability

The CMap LINCS 2020 L1000 data are publicly available at https://clue.io/data/CMap2020#LINCS2020. The processed L1000 dataset and the CRISPR knock-out dataset are available at https://github.com/Bunnybeibei/GexMolGen. The sci-Plex3 perturbation dataset is available at https://doi.org/10.6084/m9.figshare.22122701. The single-nucleus transcriptomes of opioid overdose cases and controls are available at https://www.ncbi.nlm.nih.gov/geo/query/acc.cgi?acc=GSE240457.

## Author Contributions

S. Z. conceived the concept, designed the method and the experiments, prepared data, implemented the algorithms, performed the experiments, analyzed results, and wrote the manuscript. R. O. prepared data and performed the experiments. M. M. prepared data. L. X. conceived the concept, designed the method and the experiments, wrote the manuscript, and acquired funding.

