## Supplementary Tables and Figures for "GEM-GPT Enables Personalized Cell Type-Resolved Therapeutic Design for Systems Pharmacology"

#### 1 Supplementary Tables

**Supplementary Table 1** Ablation study of GEM-GPT. Average Morgan fingerprint-based Tanimoto similarity is reported for the full model and different ablated variants. Higher values indicate better performance.

| Model variant | Average Tanimoto Similarity (Morgan FP) ↑ |
|---|---|
| **GEM-GPT** | **0.223** |
| w/o deep fusion | 0.182 |
| w/o scGPT | 0.110 |
| w/o cell state embedding | 0.005 |
| w/ concatenation fusion | 0.197 |

**Supplementary Table 2** Comparison of Boltz-2-predicted binding probabilities between GEM-GPT and baseline methods. Values under "Mean" represent the mean Boltz-2-predicted binding probability for molecules generated by each method. $p$ values were calculated using one-sided Mann-Whitney U tests comparing the binding probability distributions of GEM-GPT with those of the corresponding baseline method.

| | GEM-GPT | MolGAN | | GxVAEs | | MolGene-E | |
|---|---|---|---|---|---|---|---|
| Target | Mean | Mean | $p$ | Mean | $p$ | Mean | $p$ |
| AKT1 | **0.136325** | 0.069340 | $2.00 \times 10^{-6}$ | 0.074712 | $3.60 \times 10^{-5}$ | 0.097212 | $1.31 \times 10^{-2}$ |
| AKT2 | **0.141942** | 0.085470 | $5.00 \times 10^{-6}$ | 0.104577 | $1.39 \times 10^{-3}$ | 0.114894 | $3.80 \times 10^{-3}$ |
| AURKB | **0.128115** | 0.049833 | $1.13 \times 10^{-9}$ | 0.069342 | $3.87 \times 10^{-5}$ | 0.095537 | $1.16 \times 10^{-2}$ |
| CTSK | **0.228675** | 0.089450 | $2.67 \times 10^{-13}$ | 0.177538 | $7.55 \times 10^{-3}$ | 0.179158 | $2.72 \times 10^{-2}$ |
| EGFR | **0.116861** | 0.061712 | $4.71 \times 10^{-8}$ | 0.070585 | $1.68 \times 10^{-5}$ | 0.099498 | $9.70 \times 10^{-2}$ |
| HDAC1 | **0.234185** | 0.092525 | $2.28 \times 10^{-14}$ | 0.185251 | $1.90 \times 10^{-3}$ | 0.171384 | $1.38 \times 10^{-3}$ |

**Supplementary Table 3** FDA-approved drugs among GEM-GPT-generated molecules identified as strong binders to HRH1 by Boltz-2. The original generated SMILES, corresponding drug names and therapeutic categories, as well as Boltz-2-predicted binding probabilities and binding affinities (pIC50), are reported.

| **GEM-GPT Generated SMILES** | **Known Name** | **Therapeutic Category** | **Binding Probability** | **Binding Affinity** |
|---|---|---|---|---|
| COc1ccc(CN(CCN(C)C)c2ccccn2)cc1 | Pyrilamine | Antihistamines | 0.99 | 8.82 |
| Clc1ccc2c(c1)CCc1cccnc1C2=C1CCNCC1 | Desloratadine | | 0.99 | 8.72 |
| CN(C)CCC(c1ccc(Cl)cc1)c1ccccn1 | Chlorpheniramine | | 0.95 | 7.97 |
| CN(C)CCN(Cc1ccccc1)c1ccccn1 | Tripelennamine | | 0.98 | 7.85 |
| CN(C)CCC(c1ccc(Br)cc1)c1ccccn1 | Brompheniramine | | 0.92 | 7.71 |
| CN(C)CCC(c1ccccc1)c1ccccn1 | Pheniramine | | 0.92 | 7.52 |
| CN(C)CCC=C1c2ccccc2COc2ccccc21 | Doxepin | Antidepressants | 0.99 | 9.31 |
| CN(C)CCC=C1c2ccccc2CCc2ccccc21 | Amitriptyline | | 0.99 | 9.24 |
| CN(C)CCCN1c2ccccc2CCc2ccccc21 | Imipramine | | 0.99 | 9.03 |
| CN(C)CCCN1c2ccccc2CCc2ccc(Cl)cc21 | Clomipramine | | 0.99 | 8.65 |
| CNCCCN1c2ccccc2CCc2ccccc21 | Desipramine | | 0.99 | 8.48 |
| CN1CCN(C2=Nc3cc(Cl)ccc3Nc3ccccc32)CC1 | Clozapine | Antipsychotics | 0.99 | 9.05 |
| CN(C)CCCN1c2ccccc2Sc2ccc(Cl)cc21 | Chlorpromazine | | 0.98 | 8.82 |
| O=C(CCCN1CCC(O)(c2ccc(Cl)cc2)CC1)c1ccc(F)cc1 | Haloperidol | | 0.94 | 7.59 |
| CN1CCN(CCCN2c3ccccc3Sc3ccc(Cl)cc32)CC1 | Prochlorperazine | Antiemetics | 0.97 | 8.20 |
| CN1CCN(C(c2ccccc2)c2ccccc2)CC1 | Cyclizine | | 0.96 | 7.30 |

**Supplementary Table 4** GEM-GPT-generated molecules predicted by Boltz-2 to bind multiple OUD-related targets. The original generated SMILES, corresponding binding targets, and Boltz-2-predicted binding probabilities and binding affinities (pIC50) are reported.

| **GEM-GPT Generated SMILES** | **Targets** | **Binding probability** | **Binding affinity** |
|---|---|---|---|
| CN1CCN(c2ccc(Nc3ncc(Cl)c(NC4CC4)n3)cc2)CC1 | FGFR1 | 0.86 | 8.07 |
| | JAK3 | 0.73 | 7.08 |
| CC(=O)c1nc(Nc2ccc(N3CCN(C)CC3)cc2)ncc1Cl | FGFR1 | 0.78 | 7.63 |
| | JAK3 | 0.74 | 6.76 |
| C=C(O)c1nc(Nc2ccc(N3CCN(C)CC3)cc2)ncc1Cl | FGFR1 | 0.77 | 7.37 |
| | JAK3 | 0.74 | 6.77 |
| CN1CCN(c2ccc(Nc3ncc(Cl)c(O)n3)cc2)CC1 | FGFR1 | 0.78 | 6.92 |
| | JAK3 | 0.76 | 6.26 |

**Supplementary Table 5** Hyperparameter configurations of GEM-GPT across experimental settings. Training and generation hyperparameters used in each experiment are reported. N/A indicates that no additional model training was performed in the corresponding setting.

| Parameter | Bulk ID cell-line | Bulk OOD cell-line | sci-Plex3 | CRISPR knock-out | OUD |
|---|---|---|---|---|---|
| Learning rate | 3e-4 | 3e-4 | 6e-5 | N/A | 3e-4 |
| Batch size | 128 | 128 | 128 | N/A | 128 |
| Max epochs | 40 | 400 | 40 | N/A | 1000 |
| Hidden layers | 12 | 12 | 12 | 12 | 12 |
| Head of attention | 8 | 8 | 8 | 8 | 8 |
| Hidden dimension | 512 | 512 | 512 | 512 | 512 |
| Beam size | 20 | 20 | 20 | 20 | 1000 |

### 2 Supplementary Figures

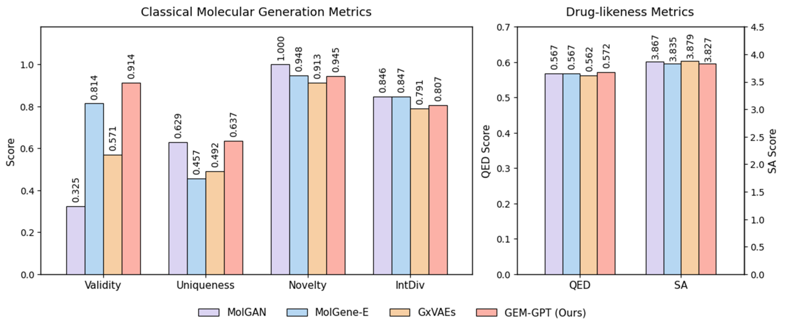


**Supplementary Fig. 1** Comparison of classical molecular generation metrics (Validity, Uniqueness, Novelty, and IntDiv) and drug-likeness metrics (QED and SA) for molecules generated from compound-induced gene expression profiles in the in-distribution (ID) cell-line setting. GexMolGen is excluded because its complete set of generated molecules is unavailable.

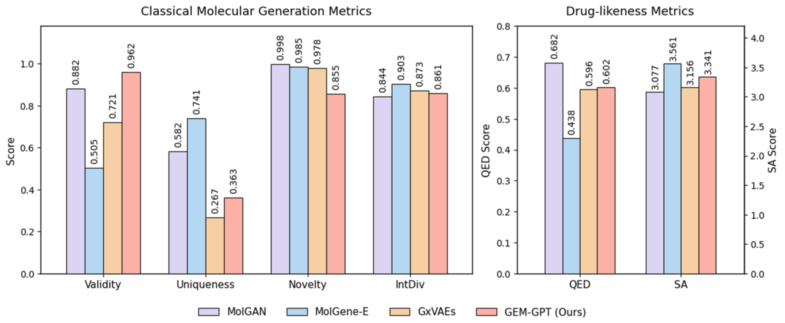


**Supplementary Fig. 2** Comparison of classical molecular generation metrics (Validity, Uniqueness, Novelty, and IntDiv) and drug-likeness metrics (QED and SA) for molecules generated from compound-induced gene expression profiles in the out-of-distribution (OOD) cell-line setting.

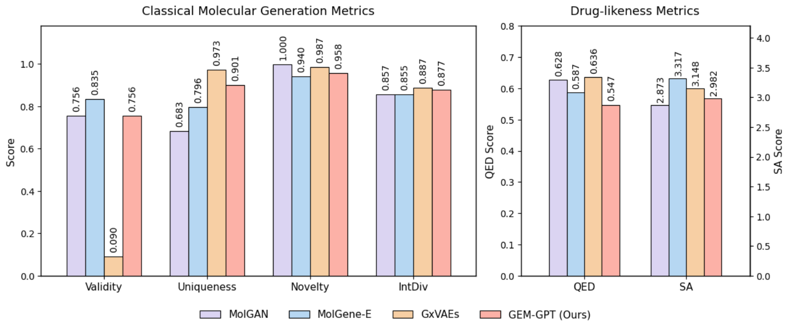


**Supplementary Fig. 3** Comparison of classical molecular generation metrics (Validity, Uniqueness, Novelty, and IntDiv) and drug-likeness metrics (QED and SA) for molecules generated from single-cell compound-induced gene expression profiles in the sci-Plex3 dataset using a compound-based split.

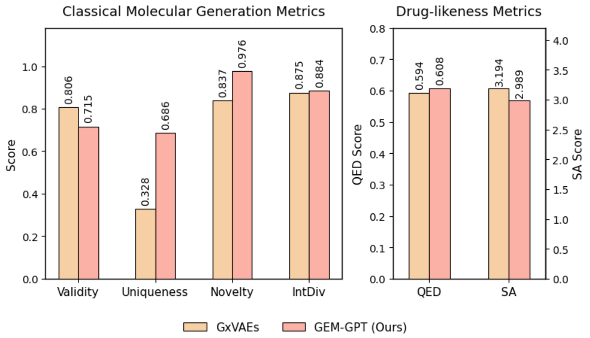


**Supplementary Fig. 4** Comparison of classical molecular generation metrics (Validity, Uniqueness, Novelty, and IntDiv) and drug-likeness metrics (QED and SA) for molecules generated from profiles derived from OUD single-nucleus RNA-seq data.

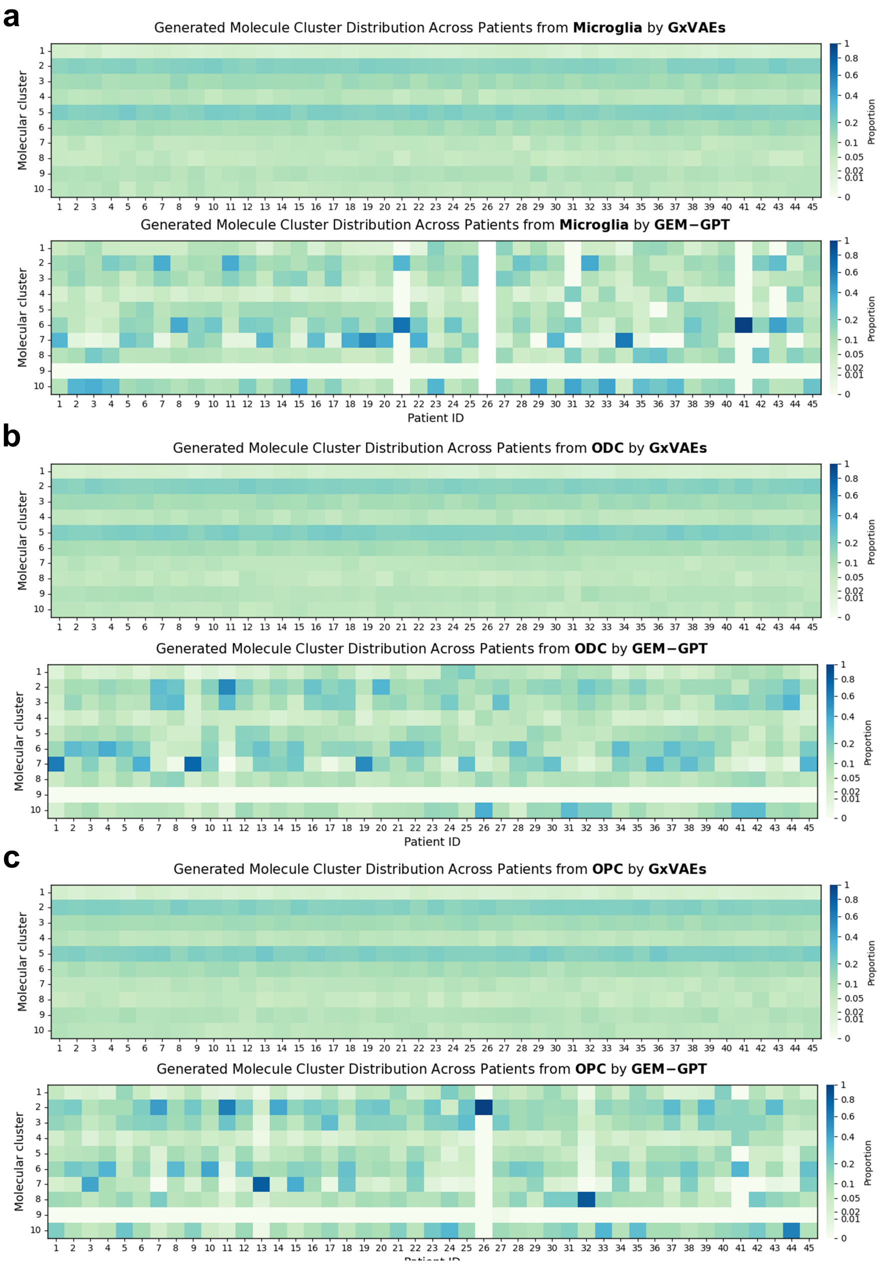


**Supplementary Fig. 5** Molecular cluster distributions of candidates generated by GxVAEs and GEM-GPT across OUD patients using cell type-specific gene expression profiles from (a) microglia, (b) ODCs, and (c) OPCs.

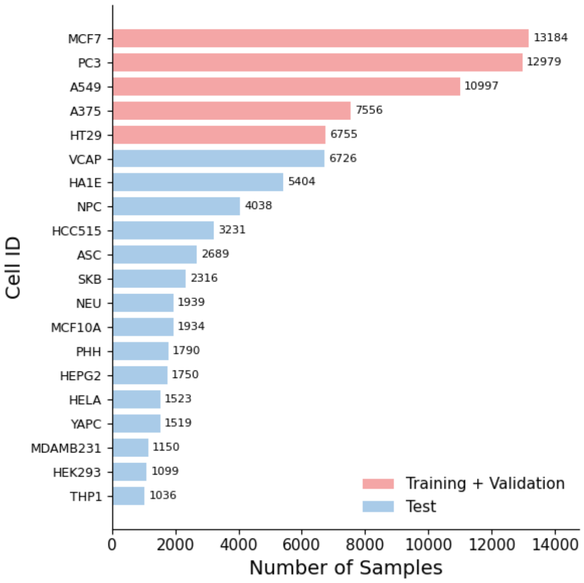


**Supplementary Fig. 6** Distribution of gene expression profile counts across cell lines in the out-of-distribution (OOD) cell-line setting.

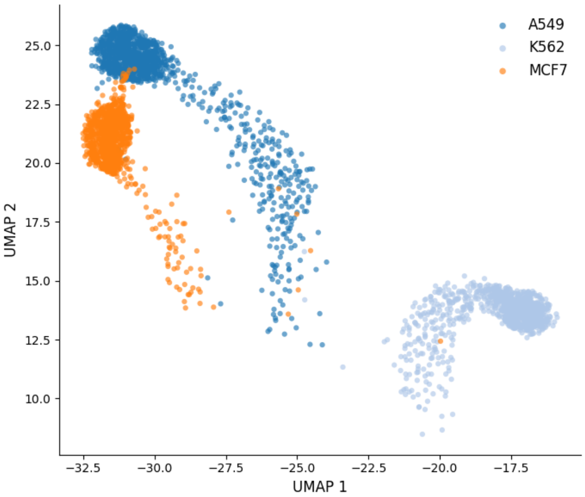


**Supplementary Fig. 7** UMAP visualization of single-cell-derived gene expression profiles from sci-Plex3, colored by cell line.

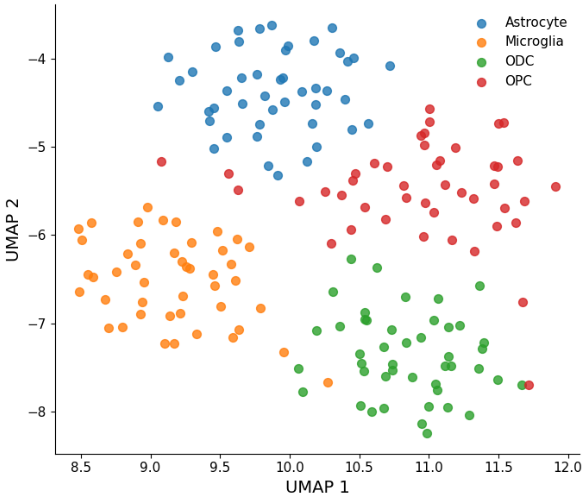


**Supplementary Fig. 8** UMAP visualization of single-nucleus-derived gene expression profiles from the OUD dataset, colored by cell type.
